# *AncestryPainter* 2.0: Visualizing ancestry composition and admixture history graph

**DOI:** 10.1101/2024.04.08.588394

**Authors:** Shuanghui Chen, Chang Lei, Xiaohan Zhao, Yuwen Pan, Dongsheng Lu, Shuhua Xu

## Abstract

To effectively depict the results of population genetics studies, it is essential to present ancestry composition and genetic distance. The growing amount of genomic data prompted us to design *AncestryPainter* 1.0, a Perl program to display the ancestry composition of numerous individuals using a rounded graph. Motivated by the requests of users in practical applications, we updated *AncestryPainter* to version 2.0 by coding in an R package and improving the layout, providing more options and compatible statistical functions for graphing. In particular, *AncestryPainter* 2.0 implements a method admixture history graph (AHG) to infer the admixture sequence of multiple ancestry populations, and allows for multiple pie charts at the center of the graph to display the ancestry composition of more than one target population. We also introduced an additional graphing module to visualize genetic distance through radial bars of varying lengths surrounding a core. Visualization functions *per se* have been enhanced in this update as well. Furthermore, *AncestryPainter* 2.0 includes two statistical modules to 1) merge ancestry proportion matrices and 2) infer admixture sequences through correlation analyses. *AncestryPainter* 2.0 is publicly available at https://github.com/Shuhua-Group/AncestryPainterV2 and https://pog.fudan.edu.cn/#/Software.

## Introduction

As the amount of sequenced and genotyped genomes grows rapidly, the analysis and depiction of the population structure and genetic affinity of a larger number of human groups have become increasingly common. The visualization of ancestry composition and genetic distance plays a crucial role in presenting the findings of population genetics studies. The conventional method of displaying ancestry composition is to align individuals in a rectangular graph, which can be challenging to print when dealing with a large number of individuals. To address the aforementioned issue, a computational program named *AncestryPainter* was thereby developed using a circular graph to display ancestry composition. Moreover, version 1.0 of *AncestryPainter* can categorize input populations based on their representative ancestry and automatically sort populations and individuals according to their ancestry proportion. Alternatively, users can specify the population order by themselves. Users can also customize the population order and modify graphic features such as ancestry colors.

Although *AncestryPainter* 1.0 has been applied to many population genetics studies, its limitations have hindered broader application. *AncestryPainter* 1.0 was written mainly in Perl but generates an R script to plot figures. The code structure inhibits users from conveniently modifying parameters when calling plotting functions within the R environment. The limited aesthetic parameters and monotonous layout, which only allows a single pie chart to highlight the ancestry of a specific population or individual at the center of the plot, further restrict its use. In addition, no statistical function compatible with plotting functions (e.g., the clustering function in the R package “*pheatmap*”) is implemented in *AncestryPainter*.

In this study, we developed version 2.0 of *AncestryPainter* using R language. This updated version retains most of the previous features while offering multiple layout styles for targets in a circular graph. Using the same basic plotting functions, we further designed a plot to display the genetic distance, in which the bars indicating genetic distance are arranged radially around the central target. To enhance the visual attraction of the plot, *AncestryPainter* 2.0 provides a variety of aesthetic parameters. Moreover, we implemented statistical functions to merge ancestry composition and infer the admixture topology in this package. Our R package aims to improve the visualization and processing of the results in population genetics studies.

## Methods

### Example dataset

To illustrate the utilities of *AncestryPainter* 2.0, we used the genome-wide SNPs of 2422 modern human individuals in the Human Origins dataset(Lazaridis, et al. 2014) and 7 Kyrgyz individuals from the Estonian Biocentre Human Genome Diversity Panel (EGDP) (Pagani, et al. 2016) to generate the example data. We converged the Human Origins and EGDP data by bcftools (Danecek, et al. 2021) and performed ADMIXTURE(Alexander, et al. 2009) to estimate the ancestry makeup of the individuals for 10 repeats, specifying the ancestry component number (*K*) as 8. In addition, we ran an in-house Python script to calculate the genetic distance (*F*_ST_)(Weir and Cockerham 1984) between populations.

### Using sectors for visualization

The graphic functions of *AncestryPainter* 2.0 are composed primarily based on the R package “graphics”. The sectors visualizing ancestry proportion or genetic distance are plotted by the function “polygon” in “graphics”. The coordinates of sectors on the canvas depend on 1) the order of the ancestry component indicated in the input data and 2) the initial plotting position. The sector size correlates with the quantity of ancestry proportion or genetic distance. In addition, we utilize other functions in the “graphics” package, such as “text” and “arrows” to annotate sectors.

### Merging ancestry proportion matrices

This section is translated from the in-house Python script authored by Pan, et al (Pan, et al. 2022) (https://github.com/Shuhua-Group/ADMIXTURE.merge). This function merges the ancestry proportion matrices (called “target matrices”) estimated by software such as ADMIXTURE with the same dataset and the same ancestry component number (*K*). This function calculates and compares the correlation (measured by Pearson coefficient) between one ancestry component in a user-defined reference matrix (i.e., a reference component) and each of the ancestry components in the matrices to be merged (i.e., target components), and then matches the reference component with the target of the highest correlation coefficient. The function counts the number of target components matched with each reference component, and calculates the supporting ratio of all ancestry components in a reference matrix. The supporting ratio is defined as the ratio of the matched target component number to the total number of target matrices. In the merged matrix, the proportion of an ancestry component for each individual is the average of a group of matched ancestry components. A target matrix with all ancestry components matching those of the reference is defined as a consensus matrix “supporting” the reference, otherwise, it is regarded as a “conflicted” one. A larger number of consensus matrices indicates the reliability of the reference and vice versa.

### Inferring admixture sequence using Admixture History Graph

Pugach, et al (Pugach, et al. 2016) have introduced a method, Admixture History Graph (AHG), to infer the admixture sequence of multiple ancestry populations by calculating correlation coefficients between ancestry components, based on the idea that the admixture proportion of two previously admixed ancestries and that of a third ancestry would be independent in subsequent admixture events. Specifically, in the AHG test, the correlation efficiency is estimated with the covariance of 1) the ratio of the admixture proportion of two random-picking ancestry components and 2) the admixture proportion of the third ancestry component. For example, an already admixed population with two different ancestry components A and B meets with another episode of admixture bringing into this population a new ancestry component C, and the arrays of ancestry proportion of individuals in populations A, B, and C are available. The correlation coefficient can be calculated as follows:

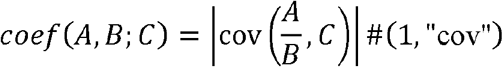

This coefficient is expected to be zero. Practically, the admixture topology with the lowest corresponding correlation coefficient among the three combinations, i.e., *coef* (*A,B;C*), *coef* (*A,C;B*), and *coef* (*B,C;A*)can be inferred as the best-fit. The supporting ratio of each admixture topology can be estimated by using ancestry proportion arrays of randomly-picked individuals from the given population.

This metric has been modified and then applied to our previous study of the Uyghurs (Feng, et al. 2017) and the Huis (Ma, et al. 2021) in Northwestern China, in which the covariance was substituted by Pearson coefficient, because the latter can adjust the bias caused by admixture proportion differences among ancestry components:

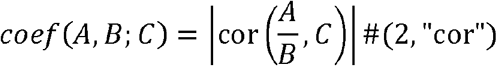

However, the correlation efficient values of the same population combination (e.g., *coef* (*A,B;C*) and *coef* (*B,A;C*)) can be distinct if the positions of ancestry components in the fraction are swapped (e.g., replacing A/B by B/A). To solve the issue, we defined a novel metric as an arithmetic mean of the two covariance or correlation values with swapping ancestry components:

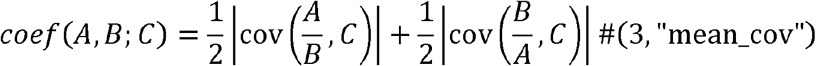

Or

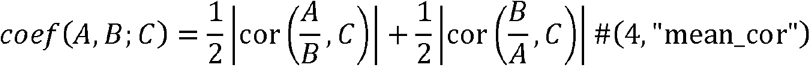

In addition, Oliveira, et al (Oliveira, et al. 2022) updated the original AHG metric by introducing logarithm-transformation, which eliminates the effect of swapping ancestry component positions.

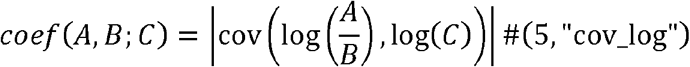

Drawing on the metrics proposed above, we could also optimize the calculation of the correlation coefficients as:

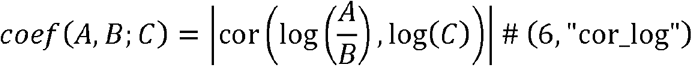

To validate the efficiency of these metrics, we examined our methods and previously-published ones via three kinds of admixture scenario models (Figure. 2). These models were established based on that proposed by Feng, et al(Feng, et al. 2017).

**Figure 1.**
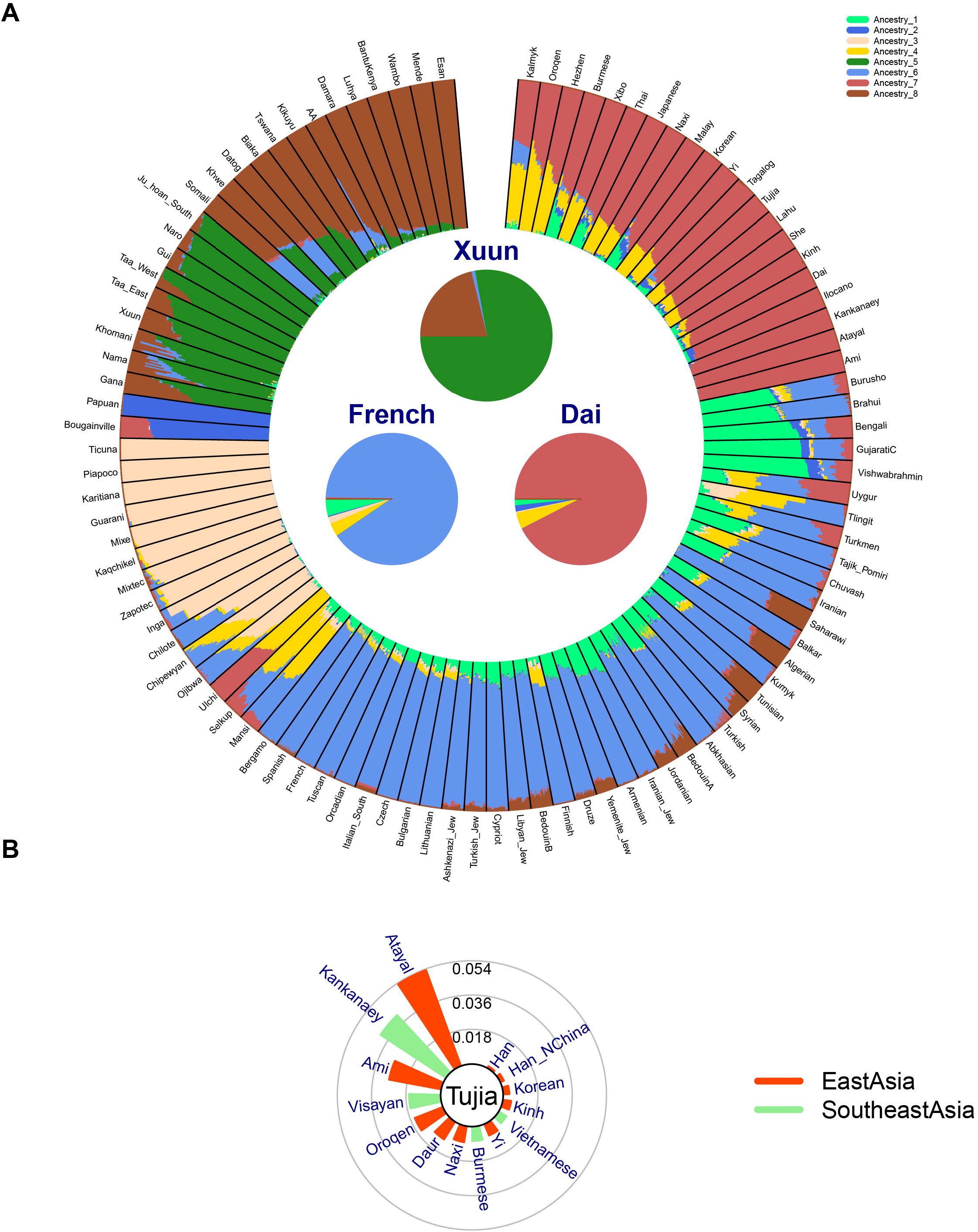
Visualization of ancestry composition and genetic distance using *AncestryPainterV2*. A) Plotted with “sectorplot” is the ancestry composition of 100 randomly-picked populations in the Human Origins Dataset given *K* = 8. The ancestry composition of three target populations (Xuun, French, and Dai) is displayed in pie charts at the center of the plot. See *Example 1* for the code. B) Plotted with “radiationplot” is the genetic distance from the Tujia group in the Human Origins Dataset to 14 randomly-picked populations from East Asia and Southeast Asia. The genetic distance is indicated by the length of the bars radially surrounding the core indicating the target. See *Example 2* for the code.

**Figure 2.**
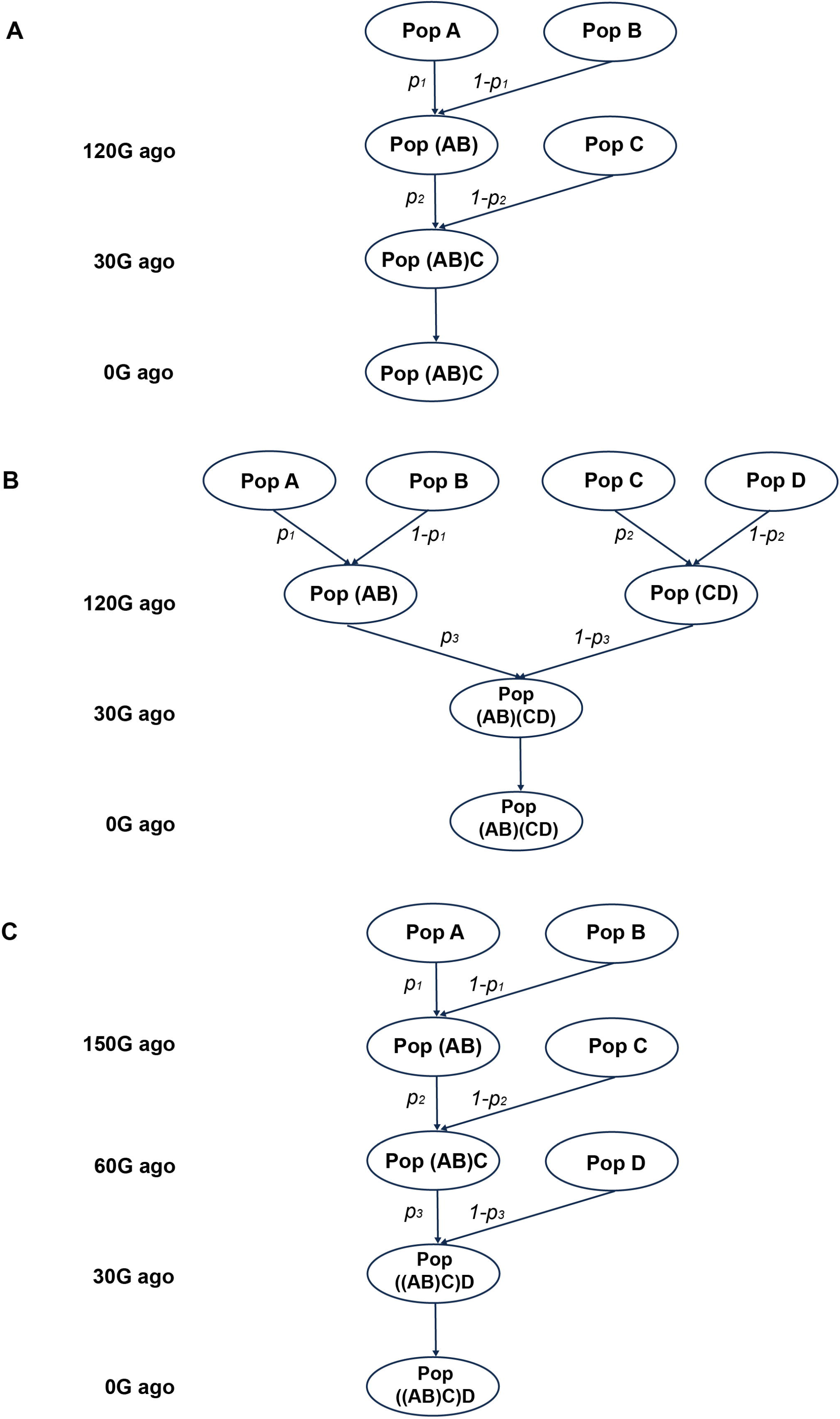
Admixture scenario models for the validation of AHG metrics. Simulated populations are marked in the oval frames, and the vertical axis left to the graph shows the admixture time. Admixture proportion is marked beside the arrows indicating admixture: A) *(AB)C* scenario, in which both *p*_*1*_ and *p*_*2*_ vary from 0.1 to 0.9, with step size as 0.1; B) *((AB)(CD))* scenario, in which both *p*_*1*_ and *p*_*2*_ vary from 0.1 to 0.9, with step size as 0.1, and *p*_*3*_ varies from 0.2 to 0.5, with step size as 0.1; C) *((AB)C)D* scenario, in which *p*_*1*_ varies from 0.2 to 0.5, with step size as 0.1, and both *p*_*2*_ and *p*_*3*_ vary from 0.1 to 0.9, with step size as 0.1.

(1) (AB)C scenario: Populations A and B were initially admixed 120 generations ago to form the AB population. After 90 generations of self-evolution, the AB population then mixed with Population C 30 generations ago, leading to the formation of the initial (AB)C population. This (AB)C population underwent a further 30 generations of self-evolution to arrive at the final (AB)C population (Figure. 2A). When population A and population B admixed to form population (AB), the admixture proportion of population A was varied incrementally from 0.1 to 0.9, with a step size of 0.1. Similarly, when population (AB) was subsequently admixed with population C, the admixture proportion of population C was also varied incrementally from 0.1 to 0.9, with a step size of 0.1. Each admixture proportion scenario was simulated 100 times.
(2) ((AB)(CD)) scenario: Populations A and B underwent admixture 120 generations ago to form the composite population (AB), while concurrently, Populations C and D were mixed to form the composite population (CD). Each of these newly formed populations, (AB) and (CD), then proceeded through 90 generations of isolated evolution before admixing 30 generations ago, giving rise to the initial combined population (AB)(CD). This combined population (AB)(CD) then experienced 30 generations of self-evolution to reach its final state (Figure. 2B). During the admixture event between Populations A and B, the contribution from Population A was incrementally set from 0.1 to 0.9 in steps of 0.1. Similarly, for the admixture between Populations C and D, Population C’s contribution was also incrementally set from 0.1 to 0.9 in steps of 0.1. During the admixture of composite Populations (AB) and (CD), the proportion of (AB) was set at 0.2, 0.3, 0.4, and 0.5. Each admixture proportion scenario was simulated 100 times.
(3) ((AB)C)D scenario: Populations A and B initially admixed 150 generations ago to form the composite (AB) population. This AB population then underwent 90 generations of independent evolution before engaging in admixture with Population C 60 generations ago, culminating in the formation of the (AB)C population. After a further 30 generations of self-evolution, this (AB)C population then admixed with Population D to form the ((AB)C)D population 30 generations ago. The ((AB)C)D population continued to evolve independently for an additional 30 generations to achieve its final genetic composition (Figure. 2C). During the admixture event between Populations A and B, the admixture proportions from Population A were specifically set at 0.2, 0.3, 0.4, and 0.5. When Population (AB) admixed with Population C, the admixture proportion from Population C varied sequentially from 0.1 to 0.9, with a step increment of 0.1. Subsequently, for the admixture event between Population (AB)C and Population D, Population D’s admixture proportion also ranged sequentially from 0.1 to 0.9, with the same step increment of 0.1. Each admixture proportion scenario was simulated 100 times.

Populations in the three kinds of admixture scenario models were simulated by AdmixSim2(Zhang, Liu, et al. 2021), an individual-based forward-time simulation tool that can flexibly and efficiently simulate population genomics data under complex evolutionary scenarios. For all 3 scenarios, the populations A, B, C, and D were randomly generated in accordance with the AdmixSim2 manual (https://github.com/Shuhua-Group/AdmixSim2/tree/master), without involving specific population information.

During the simulation, we maintained a constant sample size of 5,000 individuals per population per generation. Due to factors such as genetic drift, some ancestral components may be represented at very low frequencies (below 1e-6) in the outcomes. These minor ancestral components are then set to a threshold value of 1e-6, and the proportional frequencies of the remaining ancestral components are accordingly adjusted to ensure that the sum of all ancestral component proportions equals 1. In addition, we forced the proportion of each ancestry component in the ultimate admixed population to be greater than 0.

Further, we generated simulated data within the (AB)C scenario to evaluate the impact of varying sample sizes on algorithmic performance. Here, the metric employed was “mean_cor’’. When population A and population B were mixed to form population (AB), the admixture proportion of population A was varied incrementally from 0.1 to 0.5, with a step size of 0.1. Similarly, when population (AB) was subsequently admixed with population C, the admixture proportion of population C also varied incrementally from 0.1 to 0.9, with a step size of 0.1. Each admixture proportion scenario was simulated 100 times. After the simulation data were prepared, we sequentially sampled 25, 50, 75, and 100 individuals from the admixed population to assess the effect of sample size on the efficacy of the algorithm. We documented the number of instances in which the algorithm accurately inferred the correct admixture model (accuracy). It turned out that a greater sampling size resulted in higher accuracy of inference and all the methods obtained the highest accuracy when the sapling size was 100 (Supplementary Table 1).

## Description

### Visualization of ancestry makeup and genetic distance

*AncestryPainter* 2.0 implements a “sectorplot” to visualize the ancestry composition of multiple populations. The users of our software have to provide an ancestry matrix with rows as individuals and columns as ancestry proportion, along with the annotation including individual ID and group ID. Users can specify the color code of ancestry components and the population order. If not, the colors of ancestry will be randomly generated, and the populations will be categorized into *K* (ancestry component number, see Methods) groups and then sorted according to their representative ancestry (i.e., the ancestry accounting for the largest proportion in this population), similar to what is done in *AncestryPainter* 1.0 (Feng, et al. 2018).

An important function and new feature of our software is to display the ancestry composition of multiple target population(s) using pie charts in the center of the plot. In contrast to the *AncestryPainter* 1.0 which allows only one pie chart indicating one target population in the center, the newly-developed version 2.0 supports multiple target pie charts. This feature is inspired by some users of *AncestryPainter* 1.0 (Sala, et al. 2019; Khan and Khan 2021; Ma, et al. 2021; Zhang, et al. 2021). The positions of the target pie charts can be adjusted via the arguments defined as 1) the distance between the centers of the target pie charts and the plot; 2) the angle between the line from the center of the plot to the center of the target pie chart and the right horizontal axis of the plot.

Moreover, we designed some optional graphing elements and features to help annotate or beautify the plot. Users can add arrows from the sector indicating one population to the corresponding target pie chart, or legends that display the color code and names of the ancestry components. Users can also modify the font, size, and color of the target labels, the position of the legend, etc. For the output figures, we removed the option in version 1.0 to output graphs in “.pdf” or “.png” format directly. Instead, users can output graphs using internal R functions “pdf” “png”, etc. Another graphing function implemented in *AncestryPainter* 2.0, “radiationplot” can be used to visualize the genetic distance from one target population to another population. The plotting pattern was first present in a publication on the ancestral origin of Tibetans (Lu, et al. 2016). The required input of this plot is a four-column matrix containing information on populations, regions, genetic differences, and color codes. This plot includes a core indicating the target population surrounded radially by the sectors showing the genetic distance, with outer rings displaying the value range. The number and numeric range of outer rings can be modified by users as well. Similar to “sectorplot”, the sectors around the core can be automatically sorted according to their values. Moreover, “radiationplot” supports aesthetics and annotation such as text size/font and legends.

### Merging ancestry proportion matrices

A statistical function compatible with graphing functions is also introduced into *AncestryPainter* 2.0 for merging multiple ancestry proportion matrices estimated with the same dataset and the same ancestry component number (*K*) to obtain the averaged ancestry proportion for each individual. The required input can be the file names of the ancestry proportion data frames or the data frames directly. Each data frame contains (2 + *K*) columns, including two columns of individual and population annotation and an ancestry proportion matrix of *K* ancestry components. Users can assign any one of the inputs as the reference matrix for merging. The function “ancmerge” outputs an *R* list including 1) a merged ancestry proportion matrix with annotation, 2) a data frame showing the representative group (with the largest ancestry proportion of the corresponding component) and the supporting ratio of each ancestry component, and two vectors showing the matrices 3) conformed and 4) conflicted with the reference.

### Inferring admixture topology

We validated the efficiency of different variations of AHG by three admixture scenario models (Fig. 3; Methods). For the “(AB)C” model, given initial admixture proportion of A and C varying from 0.1 to 0.9, the metric “cov” obtained accuracy greater than 0.8 only if the initial proportion of C was not more than 0.6, and the distribution of accuracy values was asymmetric, indicating low robustness of this method. Similarly, the metric “mean_cov” showed a weakness with an extreme ratio of A (0.1 or 0.9). Compared to metrics “cov” and “mean_cov”, the other four methods (“cor” “mean_cor” “cov_log” and “cor_log”) showed better performance, while “cor” could obtain low accuracy (< 0.8) if the proportion of C was less than 0.7. In addition, “mean_cor” and “cor_log” had higher accuracy (> 0.6) than other metrics (Fig. 3A; Supplementary Table S2).

**Figure 3.**
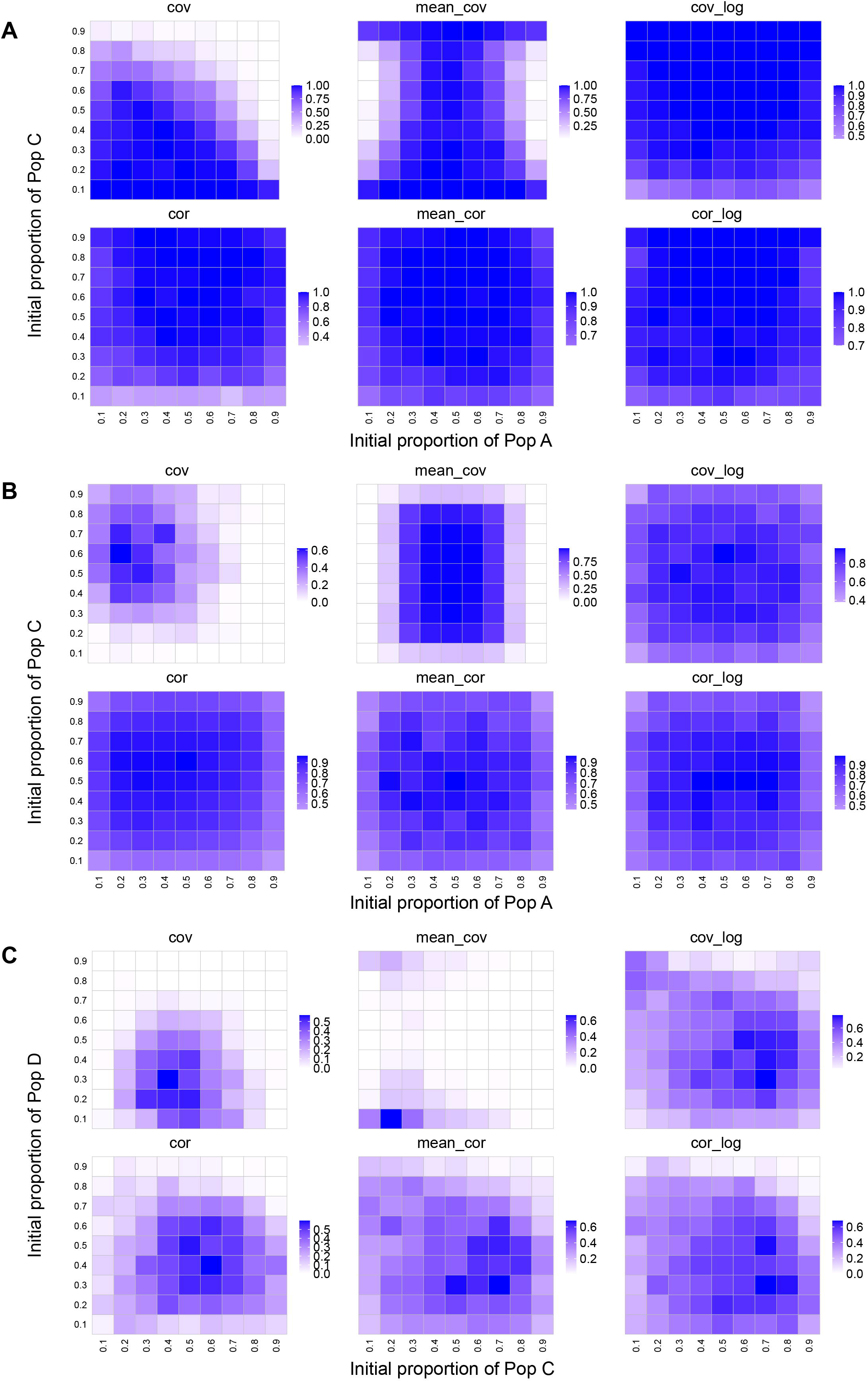
The accuracy of six AHG metrics on three admixture scenario models. The heatmaps show the accuracy of six AHG metrics on three admixture scenario models: A) (AB)C scenario; B) ((AB)(CD)) scenario; C) ((AB)C)D scenario. The accuracy value is indicated by the color gradient in the legend (high: blue; low: white). The initial proportion of the admixed population (AB) in B) was specified as 0.2. The initial proportion of the population A in C) was specified as 0.2.

When we validated efficiency of AHG metrics on the (AB)(CD) model, to make the differences of metrics more prominent, we chose a relatively biased proportion (0.2) of the admixed population (AB) (Fig. 3A), while the initial proportion of A and C varied from 0.1 to 0.9 (step size: 0.1). The metric “cov” showed the worst performance, and it was possible that “mean_cov” obtained a very low accuracy when A had an extremely low or high initial proportion (0.1 or 0.9). The rest of the metrics showed a better performance while a small proportion of A or C (0.1 or 0.9) could also reduce the accuracy. Among these metrics, “cor”, “mean_cor” and “cor_log” had relatively higher accuracy, ranging from 0.44 to 1, while the accuracy of “cov_log” might drop down to less than 0.4 (Fig. 3B; Supplementary Table S3).

For the ((AB)C)D model, we specified the initial proportion of A as 0.2, with C and D ranging from 0.1 to 0.9. Similar to the results of the (AB)(CD) model, “cov_log” “cor”“mean_cor” and “cor_log” outperformed “cov” and “mean_cov”. Moreover, “cov_log” and “cor_log” had higher median accuracy (>0.4) than “cor” (0.22) and “mean_cor” (0.38), indicating that it was more likely to obtain higher accurate admixture topology with “cov_log” and “cor_log” metrics (Fig. 3C; Supplementary Table S4).

Overall, the metric “cor_log” showed the best performance among all metrics. We further evaluated its robustness with varying (AB) and A proportion (0.3, 0.4, 0.5) in the (AB)(CD) model and the ((AB)C)D model, respectively (Fig.4; Supplementary Table S5-S6). It turned out that “cor_log” obtained an accuracy greater than 0.7 in most of the instances for the (AB)(CD) model (Fig. 4A; Supplementary Table S5). For the((AB)C)D model, if the extremely biased instances (the proportion of C or D = 0.1; the proportion of C or D = 0.9) were not taken into account, 90% of the “cor_log” results were greater than 0.5 (Fig. 4B; Supplementary Table S6). Given that admixture scenario similar to “(AB)(CD)” was more prevalent than “((AB)C)D” in previous studies(Feng, et al. 2017; Ma, et al. 2021), the lower performance of “cor_log” on the “((AB)C)D” scenario model might not hinder the application of “cor_log” in population admixture study. Therefore, we employed “cor_log” as the AHG metric of *AncestryPainter* 2.0.

**Figure 4.**
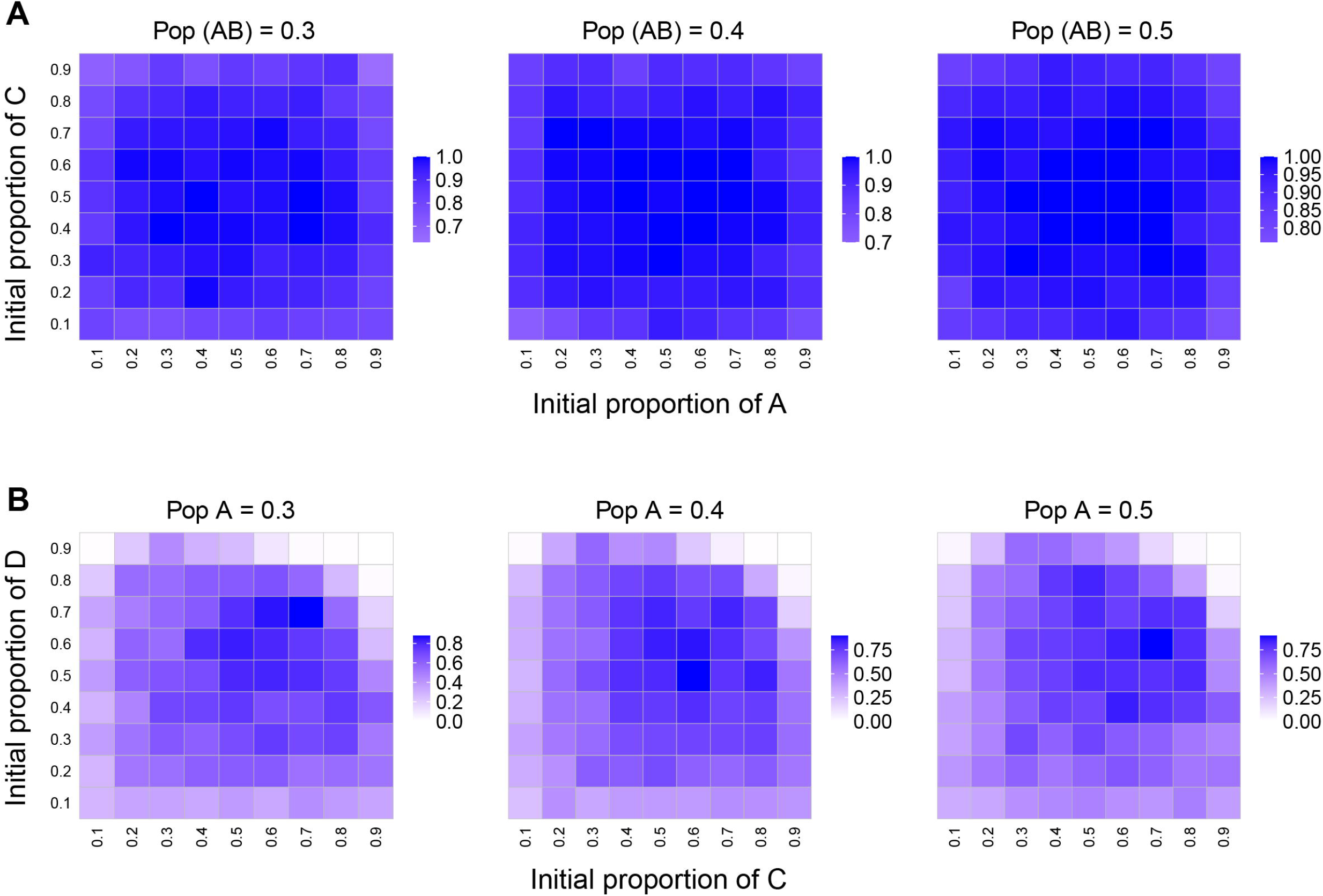
The accuracy of “cor_log” metric on two admixture scenario models. The heatmaps show the accuracy of the “cor_log” metric on A) ((AB)(CD)) admixture scenario model and B) ((AB)C)D admixture scenario model. The accuracy is indicated by the color gradient in the legend (high: blue; low: white) The initial proportion of population (AB) or population A is marked on the top of each heatmap.

## Code examples

Here are shown some examples of the usage of *AncestryPainter* 2.0. For full instructions and example data, please refer to the manual online (Data and code availability).

### Example 1: visualization for ancestry composition by sectorplot

In this example, we first read a series of input files into variables in the R environment, including the ancestry proportion matrix (“exp_q”), the individual and population annotation (“exp_ind”), the previously sorted population order (‘exp_order’) and the color code of the ancestry components (“exp_cols”).

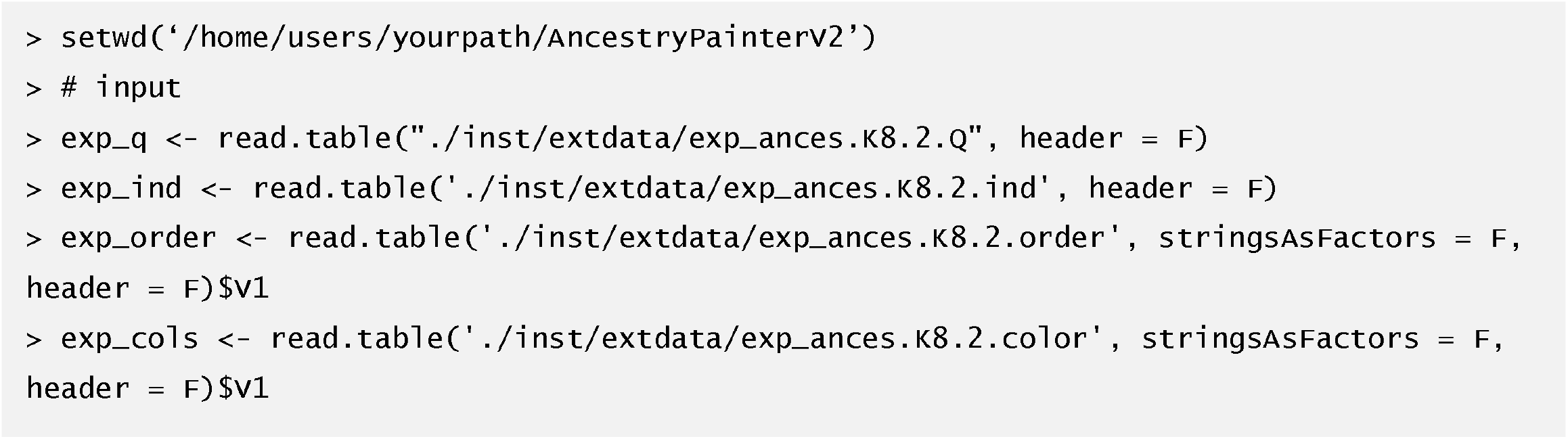

Next, these variables are passed to the “sectorplot” function with the target specified as the populations “Xuun”, “French” and “Dai”. The layout of the target pie charts is determined by the parameters “tarang1” and “tarang2”, which means the angle of the center of the pie chart in the polar coordinate system with the plot center as the origin. In addition, the flag “legend_mode” is specified as TRUE to show the legend of the ancestry components. The output is written in a PDF file and is illustrated in Fig. 1A.

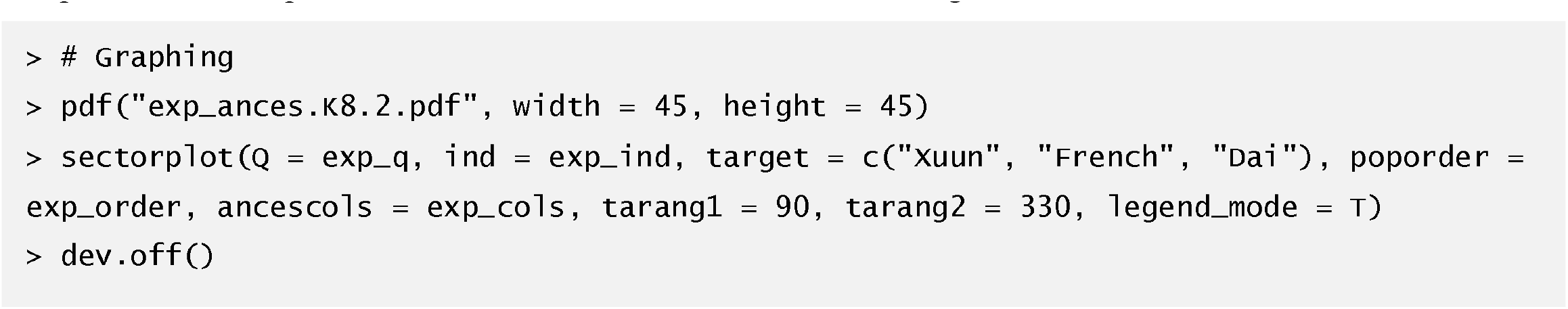

### Example2: visualization for genetic distance

Below is shown an example to use our R package to visualize the genetic distance from a target “Tujia” to reference populations, a tab-separated file is read into an R data frame and passed to the “data” parameter of “radiationplot”. The population label which is to be shown in the core of the plot should be passed to the “target” parameter. The “sorting” flag controls whether to sort the bars indicating genetic distance. Regarding the outer rings marking the genetic distance values, the number of the rings and the decimal of the values on the rings are controlled by parameters “num” and “digits”, respectively. Similar to “sectorplot”, users can decide whether to show the legend (“legend_mode”) and adjust the legend position (“legend.pos”). Moreover, the text size in the plot (e.g., “ring.text.cex”) can be modified as well. The output is illustrated in Fig. 1B.

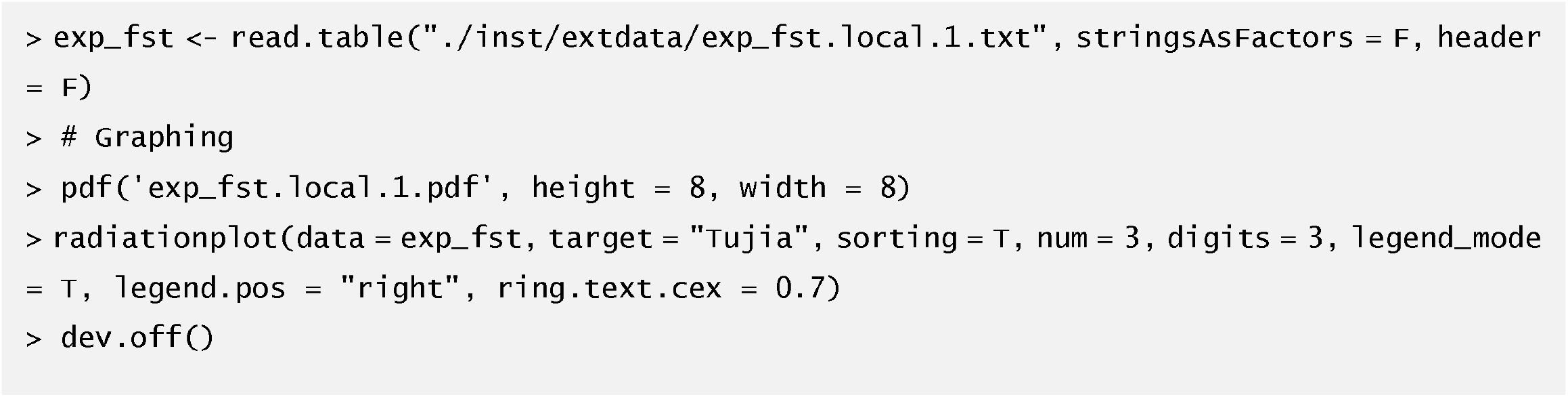

### Example3: Ancestry composition merging

In this example, we use an internal R function “list.files” to read the names of ancestry proportion files into a vector, which is then passed to the parameter “tar_anc_filelist” of the function “ancmerge” with “K” specified as 8. The reference ancestry proportion matrix is specified as the first one of the matrices. The function prints the time and working directory, and prints “Done.” after all of the comparing and merging processes are completed.

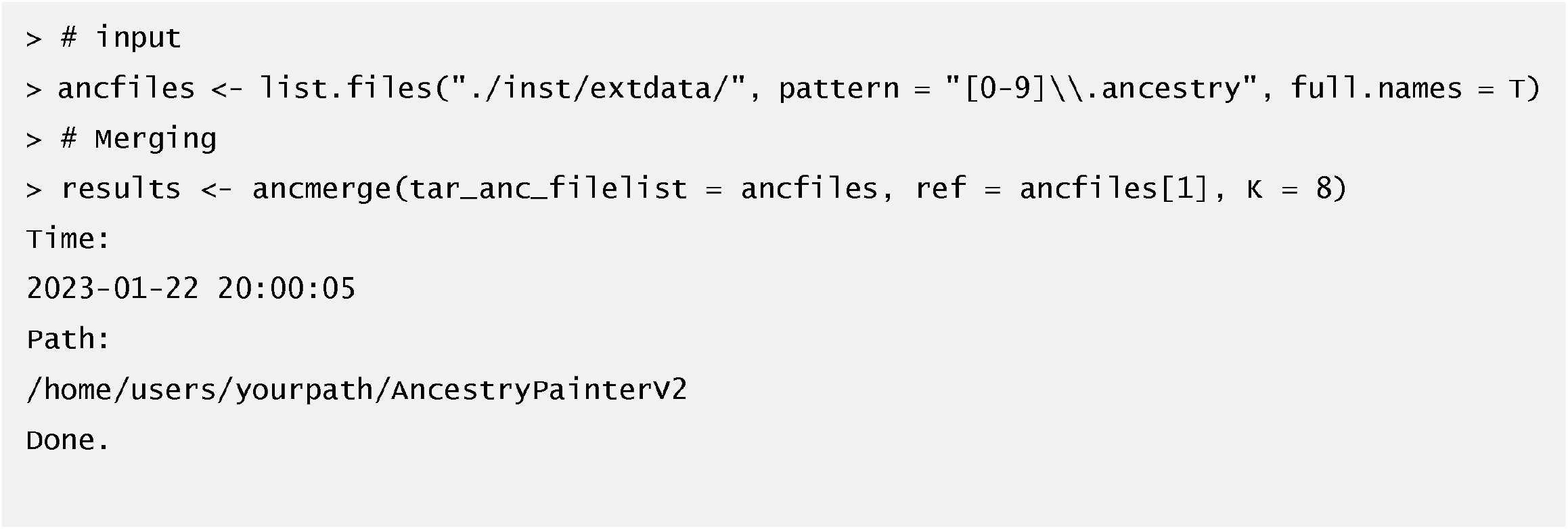

Users can view the output merged ancestry proportion matrix (“results$merged_ancesrty”) and the supporting ratio of the representative populations for ancestry components (“results$sipporting_ratio”).

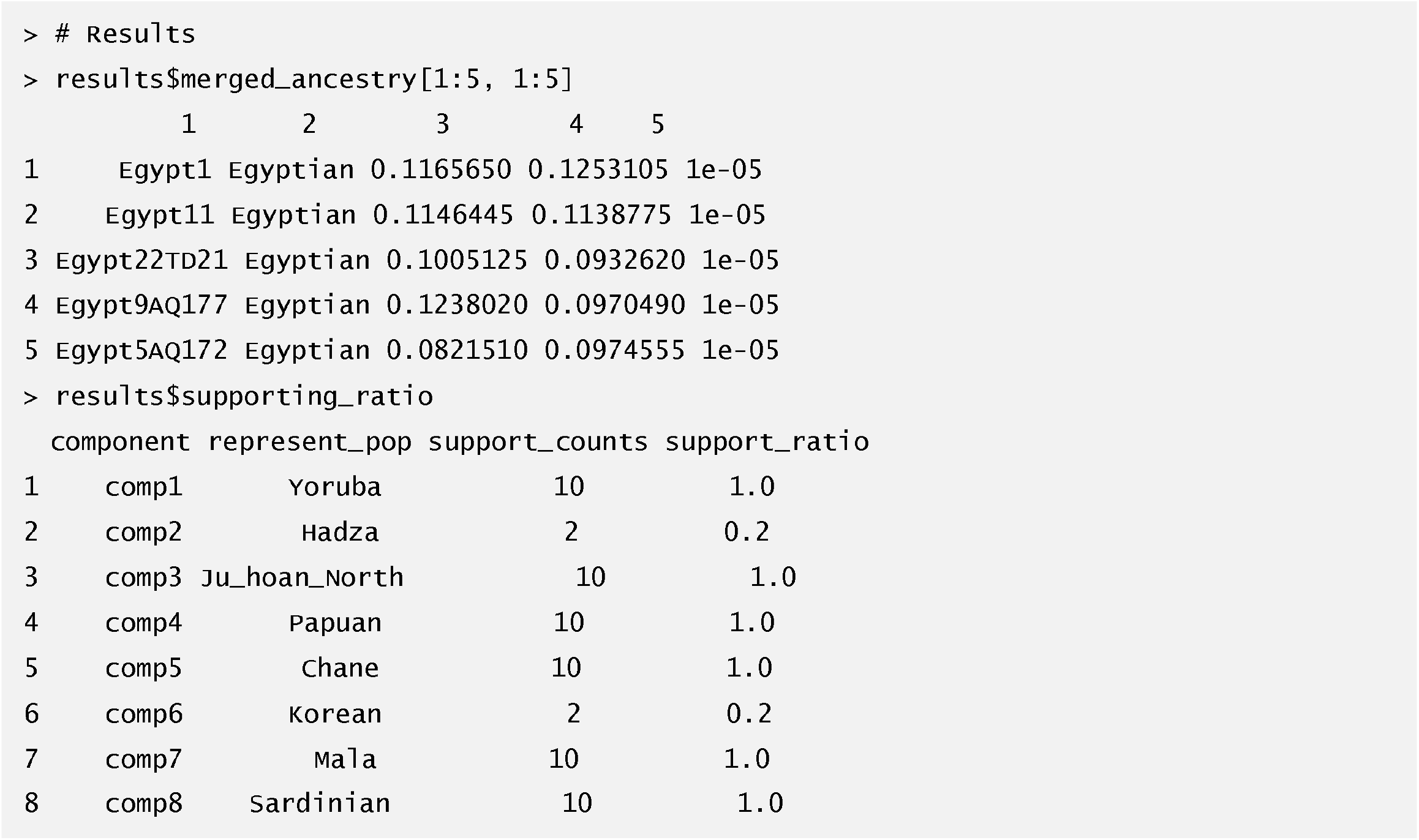

### Example4: Admixture history topology

We included the whole procedure of AHG, combination of ancestry components, sampling of individuals, and calculation of correlation coefficients in one function “ahg”. The output of this function contains 1) a data frame of all ancestry component combinations and all the possible admixture topologies together with the corresponding supporting number over all bootstrap runs; 2) the correlation coefficient of admixture topologies in each bootstrap run. Bootstrap number and sampling size can be specified via parameters “times” and “num”, respectively. It is recommended to sample as many individuals as possible to raise the accuracy of inferring the admixture topology.

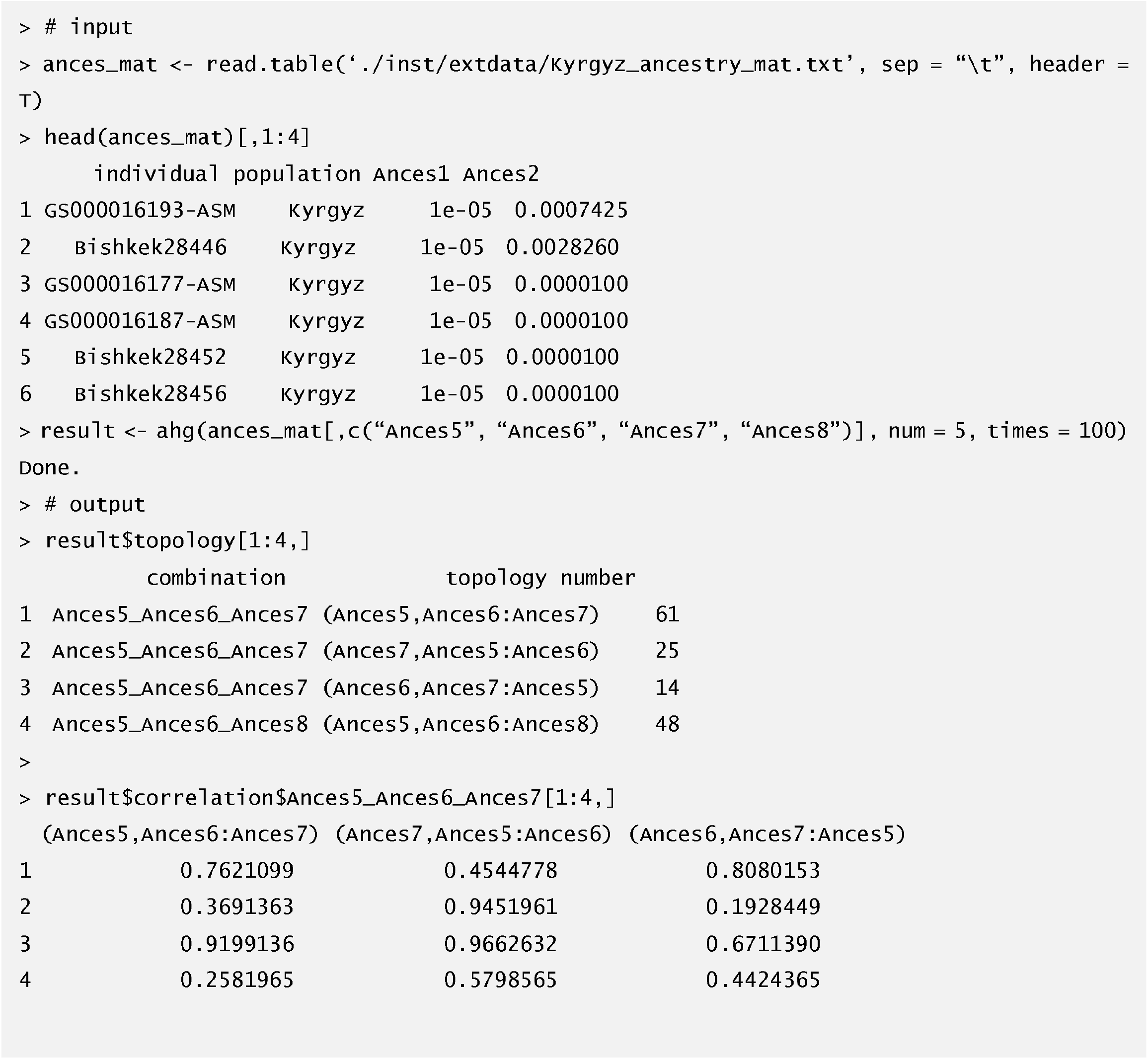

In the example above, the string “Ances5_Ances6_Ances7” indicates a combination of ancestry components “Ances5” “Ances6” and “Ances7”, and “(Ances5,Ances6:Ances7)” indicates the admixture topology in which that “Ances5” and “Ances6” admixes first and the joint ancestry admixes with “Ances7” later.

## Discussion

In this study, we developed a new version of *AncestryPainter* which can be used to illustrate the ancestry compositions and genetic distance along with statistical functions to merge multiple ancestry proportion matrices or infer admixture topology. Moreover, we introduced the AHG algorithm into *AncestryPainter* for the inference of admixture topology. We compared the accuracy of six AHG metrics on three different admixture scenario models using simulated populations. The metric “cor_log” showed an overall better performance than other metrics, and thus we implemented this metric in the AHG function.

The AHG method is easy to operate and has a high accuracy with (AB)C and (AB)(CD) admixture scenario models. However, the accuracy of all AHG metrics is low when the proportion of any ancestor is too small. It can be interpreted as the effect of genetic drift, which can be simulated by AdmixSim2. When descendants are generated, an ancestry component may be lost or drastically decreased due to genetic drift and the ancestry proportion in the descendants tends to form a truncated normal distribution with large variance, which disturbs the correlation between previously admixed ancestry components. Accordingly, all AHG metrics do not perform well in the ((AB)C)D) scenario, a continuous admixture model, which may result from the large variance of each ancestral component after admixture. The AHG accuracy for the ((AB)C)D) the scenario might grow if all four ancestral proportions have a substantial admixture proportion (Fig. 4B). Collectively, AHG can be used as a “preliminary estimate” to infer the admixture topology and have to be combined with other methods, e.g., the three-population test (*f*_3_) (Patterson, et al. 2012).

In the future, we plan to implement more functions and features, for instance, using multiple concentric circles in a single image to allow the displaying of ancestry makeup assuming different numbers of ancestry components, or annotating the subgroup information on a finer scale. Furthermore, we may introduce tree structure or network graphs to display the phylogenic relationship of populations and admixture topology.

## Data and code accessibility

Example data and source code are available on GitHub (https://github.com/Shuhua-Group/AncestryPainterV2) and HumPOG website (https://pog.fudan.edu.cn/#/Software).

## Authors’ contributions

S.X. conceived and designed the study and supervised the project. S.C., C.L., H.Z., Y.P., and L.D. contributed to the computer code. C.L. developed a key algorithm. H.Z. examined and improved the computer code. S.C. coordinated the computer coding, and packed the software. S.C. and C.L. drafted the manuscript. S.X. revised the manuscript. All authors read and approved the final manuscript.

## Acknowledgment

We thank Dr. Qidi Feng for sharing her experience on the first version of this package. This study was supported by the National Key Research and Development Program of China (No. 2023YFC2605400), the National Natural Science Foundation of China (NSFC) grants (32288101, 32030020), the Shanghai Science and Technology Commission Program (23JS1410100), the Office of Global Partnerships (Key Projects Development Fund). The computational work in this study was supported by the CFFF Computing Platform and the Human Phenome Data Center of Fudan University.

